# Distinct nociresponsive region in mouse primary somatosensory cortex

**DOI:** 10.1101/2021.04.14.439725

**Authors:** Hironobu Osaki, Moeko Kanaya, Yoshifumi Ueta, Mariko Miyata

**Affiliations:** Division of Neurophysiology, Department of Physiology, Graduate School of Medicine, Tokyo Women’s Medical University, Tokyo 162-8666, Japan

## Abstract

Nociception, somatic discriminative aspects of pain, is represented in the primary somatosensory cortex (S1), as is touch, but the separation and the interaction of the two modalities within S1 remain unclear. Here, we show the spatially-distinct tactile and nociceptive processing in the granular barrel field (BF) and the adjacent dysgranular region (Dys) in mouse S1. Simultaneous recording of the multiunit activity across subregions reveals that Dys responses are selective to noxious input whereas those of BF are to tactile input. At the single neuron level, nociceptive information is represented separately from the tactile information in Dys layer 2/3. In contrast, both modalities are converged in a layer 5 neuron in each region. Interestingly, the two modalities interfere with each other in both regions. We further demonstrate that Dys, but not BF, activity is critically involved in neuropathic pain and pain behavior, and thus provide evidence that Dys is a center specialized for nociception in S1.

## Introduction

The primary somatosensory cortex (S1) plays a central role in tactile information processing^1^. The tactile representation in S1 is orderly arranged in a somatotopic fashion^2^. On the other hand, S1 is responsible for somatic discriminative aspects of pain processing, such as the location, intensity, and quality of pain^3–10^. S1 receives thalamocortical nociceptive information^11^ and relays it to other pain-related cortical areas, such as the anterior cingulate cortex, which is responsible for the affective aspects of pain^12,13^. S1 also modulates noxious inputs via the corticotrigeminal^14^ and corticospinal^15^ pathways under both acute and chronic pain conditions. Therefore, S1 can be viewed as a network hub of pain processing and a target for interventions to control pain. However, it remains unclear how S1 processes nociceptive information and somatic tactile information distinctively.

Mouse S1 is divided into two subregions based on the cytoarchitecture: the granular region known as the barrel field (BF), which is identified by unique clusters of layer (L4) neurons, and the adjacent dysgranular region (Dys), which has poorly defined L4^6,16^. The two subregions are thought to be functionally different. For instance, BF is the center for processing tactile input from whiskers^17–19^, while Dys receives proprioceptive input by deep muscle stimulation or joint rotation^20,21^. In nociception, BF neurons in deeper layers receive noxious inputs^11,22,23^. Similarly, Dys neurons in the deeper layers respond to noxious pinching and pruriceptive inputs^24,25^. However, it is still unclear how each subregion processes nociceptive information with/without tactile information because a direct comparison between the two subregions is lacking.

Here, we found that nociceptive and tactile information is separately represented in Dys and BF, respectively, by simultaneous recording from both subregions. Dys was also predominantly activated under neuropathic pain condition. Reflecting the spatially-distinct representation of nociception, optogenetic inhibition of neuronal activity of Dys, but not BF, reduced pain behavior. Thus, we clarified a distinct functional role in nociceptive processing of Dys, which generates proper escape behavior from noxious inputs and is a potential target for pain relief.

## Results

### Nociceptive information is mainly processed in Dys

First, we sought to identify the area responding to noxious input in S1 by observing the expression of c-Fos, a neural activity marker, after capsaicin was injected into the whisker pad (Fig. 1a, b). The number of c-Fos-positive neurons increased significantly in L4 of Dys after capsaicin injection (*P* = 1.2 × 10^-5^) but not in L4 of BF (*P* = 0.97, Fig. 1b and Supplementary Table 1). The noxious stimulus-induced increase in L4 c-Fos-positive neurons was also detected in the hindpaw area of Dys when formalin was injected into the hindpaw (Extended Data Fig. 1). Thus, Dys neurons responded to noxious input in a somatotopic manner.

**Figure 1.**
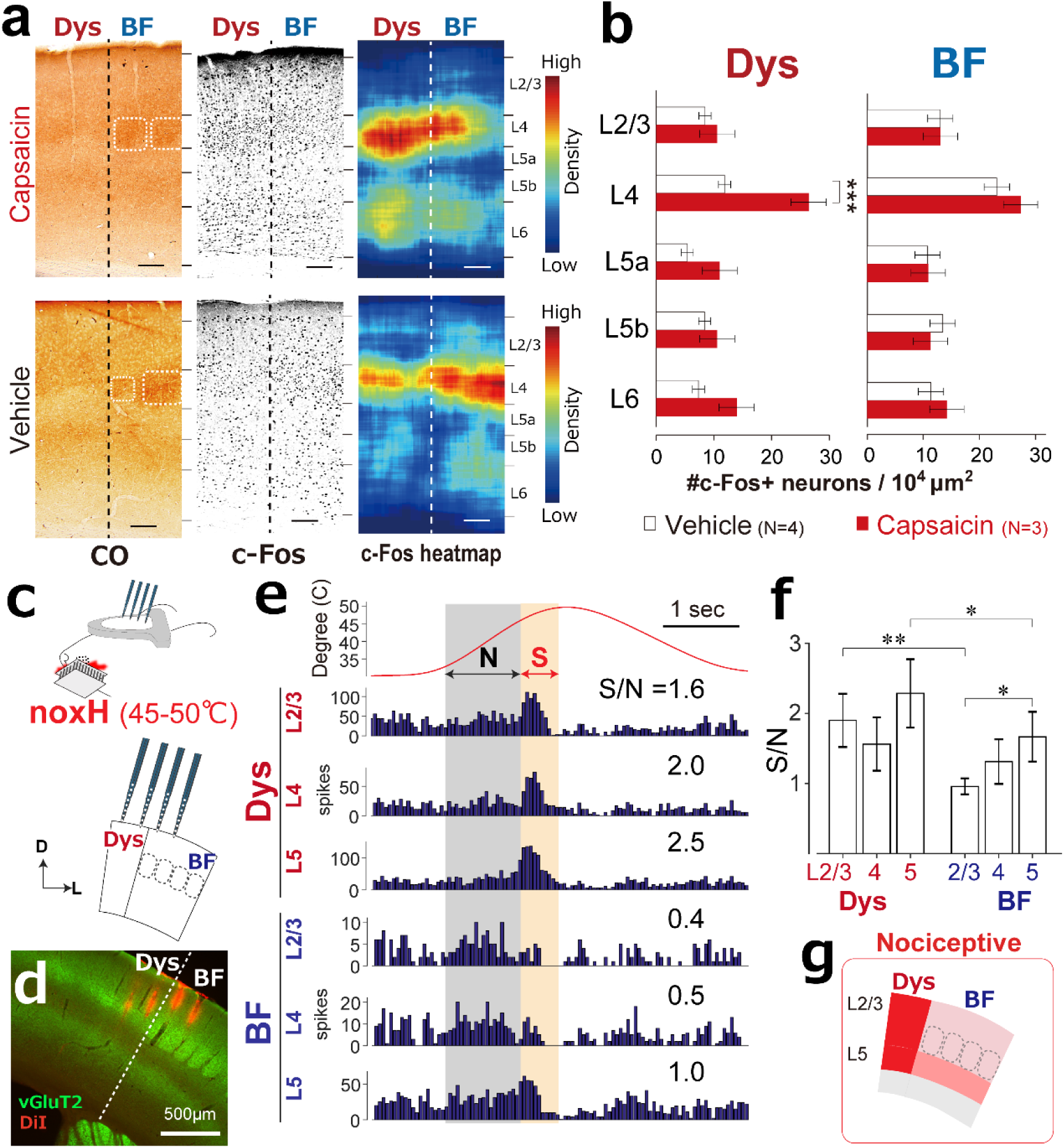
Dysgranular region in S1 responds to noxious input. **a**, Neurons in the dysgranular region (Dys) were activated by injecting capsaicin into the whisker pad. *Left*, cytochrome c oxidase (CO) staining identified the boarder of Dys and the whisker barrel field (BF). *Middle*, c-Fos immunostaining of adjacent slices is shown. *Right*, Heat maps indicate the density of c-Fos-positive neurons. Scale bars, 100 μm. **b**, Capsaicin injection into the whisker pad increased the number of c-Fos-positive neurons in Dys L4 (mean ± SEM; \*\*\**P* = 1.2 × 10^-5^, one-way ANOVA followed by Tukey–Kramer test) but not in BF L4 (*P* = 0.97). **c**, Setup for simultaneous recordings from Dys and BF after application of noxious heat stimulus (noxH) to whisker pad. **d,** Brain section showing electrode tracks (DiI, red). vGluT2 (green) staining shows the border of Dys and BF. **e,** Peristimulus time histograms of representative multiunit activity recordings to noxH in L2/3, 4, and 5 of Dys and BF. The shaded areas indicate regions for calculating ratios of signal (S) to noise (N). **f,** Statistical comparison of S/N responses to noxH. \**P* < 0.05, \*\**P* < 0.01, *n* = 8 animals; Kruskal–Wallis test followed by Dunn’s test. **g**, Summary diagram indicates that S/N for noxH was higher in Dys than BF.

Next, we compared response properties between Dys and BF, during a noxious heat stimulus (noxH; 45–50°C) applied to the whisker pad (Fig. 1c, d). We recorded multiunit activities (MUA) simultaneously from Dys and BF neurons in layer 2/3 (L2/3), L4, and layer 5 (L5) (Fig. 1e). The MUA in Dys increased in all of the recorded layers when the temperature of the Peltier device reached a noxious range (45–50°C) (Fig. 1e), while the responses differed among the layers in BF; MUA to noxH did not increase in BF L2/3 or L4 but increased slightly in L5 (Fig. 1e, bottom charts). To evaluate the selectivity to noxH, the signal-to-noise ratio (S/N; see Methods) was calculated. The response to noxH (labeled S, beige shaded region in Fig. 1e) was used as the signal, and the response to an innocuous heat range (33–45°C, labeled N, grey shaded region in Fig. 1e) was used as the noise. When comparing the S/N values for the same layers between Dys and BF(*n* = 8 animals), the S/N in Dys was significantly higher than that in BF (Fig. 1f) in L2/3 and L5 by a multiple-comparisons test (*P* = 0.9). Although L4 neurons did not show the significant difference in Figure. 1f, the comparisons of simultaneously recorded neural pairs showed that L4 neurons in Dys were significantly more selective to noxH than those in BF (Extended Data Fig. 2, *P* = 0.0056). Within BF, the S/N was significantly higher in L5 than L2/3 (*P* = 0.025, Fig. 1e). This difference between layers for noxH responses in BF was reported in the previous studies^22,23,26^. Together, the MUA analyses indicate that Dys responded more selectively to noxH than BF (Fig. 1g).

**Figure 2.**
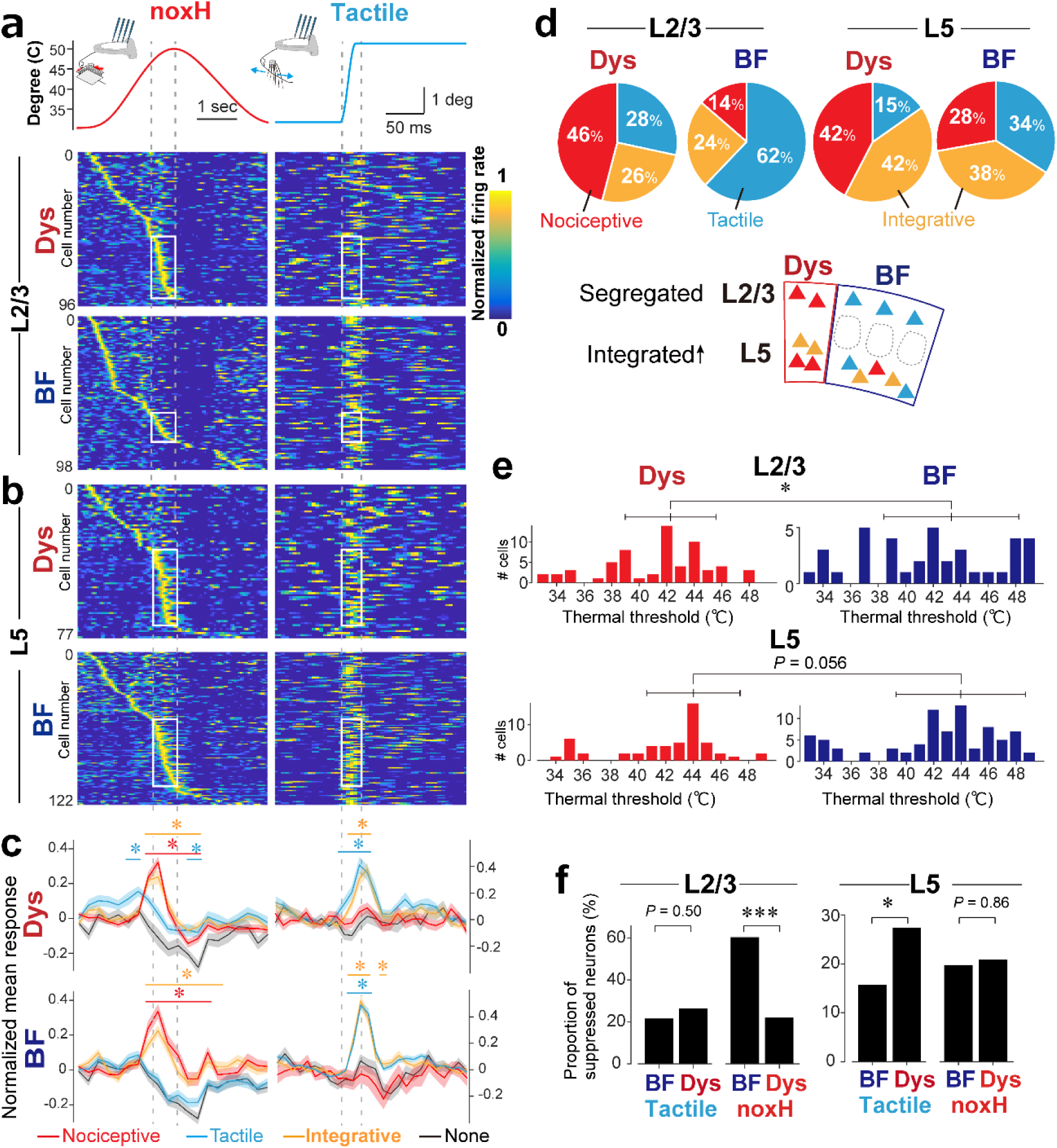
Separate processing and interaction of nociception and tactile information in S1. Peristimulus time histograms for the responses of the same L2/3 (**a**) and L5 (**b**) neurons to noxious heat (noxH) (*left*) and tactile (*right*) stimuli. 50 ms/bin for heat stimulus, 5 ms/bin for tactile stimulus. Each row was sorted by peak response time for noxH, and firing rate was normalized by peak response (boxed areas) in each row. **c**, Mean values from the histograms in panels a and b sorted according to S/Ns to noxH and tactile stimuli: nociceptive, tactile, integrative (nociceptive and tactile), and nonselective cells (none) (see Extended Data Fig. 4). \**P* < 0.05 versus nonselective. **d,** Cell type distributions in L2/3 and L5 of each area and summary diagram. **e,** Distributions of the thermal thresholds, determined from the temperature at which the neural responses reached 80% of peak spike rate. Dys neurons were tuned to noxious heat whereas BF neurons responded to various temperatures (L2/3 median ± SD: BF, 43.9°C ± 4.6°C; Dys, 42.9°C ± 3.4°C; \**P* = 0.013, two-sample Ansari–Bradley test for dispersions) (L5 median ± SD: BF, 43.9°C ± 4.7°C; Dys, 44.3°C ± 3.4°C). **f**, Proportions of neurons for which responses were suppressed (S/N < 1) or facilitated (S/N ≥ 1). \**P* = 0.048, \*\*\**P* = 5.4 × 10^-8^, 2×2 χ^2^ test between BF and Dys.

We next assessed response selectivity to tactile stimuli by comparing the S/Ns in response to whisker deflection (see Methods). Neurons in BF responded precisely to the onset of each whisker deflection, whereas those in Dys did not (Extended Data Fig. 3a, b). The S/Ns to whisker deflection were higher in BF than Dys in each layer (Extended Data Fig. 2, right, and 3c), indicating that the selectivity to tactile stimuli was higher in BF than Dys (Extended Data Fig. 3d).

### Nociceptive information is processed separately from tactile information in Dys L2/3

The results of the MUA analysis indicated that nociceptive information was mainly processed in Dys rather than BF (Fig. 1). Thus, we examined the modality specificity of single neurons in the two subregions. The peristimulus time histograms (PSTHs) of the well-isolated neurons in L2/3 simultaneously recorded from the two subregions are shown in Fig 2a; recorded neurons were sorted by the time of peak response to a heat stimulus. In Dys, the steepness of peak responses to a heat stimulus increased within the noxious heat range, indicating that many neurons responded to the noxious heat, whereas a few neurons responded in BF L2/3 (white boxes in Fig. 2a, left column). On the other hand, many BF neurons, but only a few Dys neurons, in L2/3 responded well to tactile stimuli (whisker deflections). Notably, the noxious heat-responding neurons in Dys L2/3 did not respond to the tactile stimuli (white box in Fig. 2a, right column).

In L5, the steepness of peak responses to heat stimulus increased within the noxious heat range in both regions (white boxes in Fig. 2b, left column). In contrast to that in L2/3, many nociceptive neurons in L5 of both Dys and BF responded to tactile stimuli (white boxes in Fig. 2b, right column). Thus, Dys neurons in L2/3 process mainly nociceptive information separately from tactile information, whereas neurons in L5 of both subregions tend to respond to tactile and noxious inputs.

To quantify these observations, we classified the neurons into nociceptive, tactile, integrative, and nonselective types according to S/Ns to noxH and tactile stimuli (Fig. 2c and Extended Data Fig. 4). The distributions of the S/Ns clustered according to the median S/Ns for all recorded neurons (1.81 for nociceptive and 1.46 for tactile, Extended Data Fig. 4b, d). In addition, normalized PSTHs of neurons classified according to these values represented the characteristics of each neuron group (Fig. 2c): nociceptive-type neurons responded to noxious heat but not to the tactile stimulus, tactile-type neurons responded to tactile input but not to noxious heat, and integrative-type neurons responded to both stimuli. Therefore, we used the median S/Ns as the cutoff values for classification. According to this classification, the proportion of neurons of each type in each subregion was calculated (Fig. 2d). Consistent with the population PSTH (Fig. 2a, b), in L2/3, nociceptive information was processed separately in Dys while tactile information was processed in BF. On the other hand, nociceptive and tactile information tended to be integrated at L5 neurons of both subregions (Fig. 2d, bottom).

In the population PSTHs (Fig. 2a, b), the onset of the response to heat stimuli varied across neurons, indicating that they responded to various temperatures. Therefore, we examined the thermal threshold of neurons in response to the onset of a heat stimulus (Fig. 2e). The histogram of the thermal thresholds of L2/3 neurons shows that many Dys neurons started to respond near noxious temperatures (42–44°C). By contrast, BF neurons responded to a broad range of temperatures (*P* = 0.013), suggesting that BF neurons encode cutaneous temperature^27,28^. In L5, the distribution in Dys was sharply tuned to a noxious heat range, although the difference between the distributions for Dys and BF was insignificant (*P* = 0.056). These data demonstrate a preference of Dys neurons for noxious heat.

The averaged PSTHs (Fig. 2c) show that the responses of tactile and nonselective neurons were suppressed by noxH. Because the S/Ns of these neurons were <1, we estimated the proportion of neurons suppressed by noxH in each region. In L2/3 of BF, 60% of neurons were suppressed by noxH. This proportion is significantly larger than that in Dys (22%, *P* = 5.4 × 10^-8^, Fig. 2f). In L5, on the other hand, the proportion of neurons that were suppressed by tactile stimuli was larger in Dys than BF (*P* = 0.048, Fig. 2f). These data may provide the neural mechanism underlying the interference between touch and pain^29–31^.

In summary, the data show that nociceptive information is processed separately from tactile information in Dys. The majority of nociresponsive neurons in BF were the integrative type in both L2/3 and L5. The difference in thermal thresholds indicates that Dys processes noxious heat input, and BF is responsible for temperature coding. Furthermore, the modalities interacted with each other in a way that each suppresses the other’s region.

### Dys is involved in neuropathic pain

S1 is also activated under chonic pain condition^12,13,32,33^. Thus, we next investigated how cortical representation would shift from tactile to nociceptive information in S1 during the developing of tactile allodynia induced by nerve injury. We ligated the infraorbital nerve (ION) as a trigeminal neuralgia model. For ION ligation, we used an absorbable surgical thread (see Methods), which enabled us to observe the S1 regions responding to tactile stimulus both during nerve injury and after recovery (Fig. 3a, b). Intrinsic signals in BF induced by whisker stimulation were observed before ligation (Fig. 3b, left). At postoperative day 7 (POD 7), when the tensile strength of the surgical thread is reduced to ∼50%, the signal in BF disappeared but was enhanced in the adjacent region (Fig. 3b, middle). Subsequent histological analyses confirmed that the adjacent region was Dys (Fig. 3c and Extended Data Fig. 5). At POD 21, during recovery from the ligation when the tensile strength is reduced to 0% of the maximum strength, the signal in BF reappeared (Fig. 3b, right). In population analyses, ION ligation increased the signal in Dys (Fig. 3d). Consistent with this, the number of c-Fos-positive neurons in Dys was significantly increased after ION ligation (Extended Data Fig. 6a–c). Moreover, mechanical allodynia at the whisker pad was also observed (Extended Data Fig. 6d, e), suggesting that Dys may be related to the peripheral nerve injury-induced neuropathic pain. The activated region changed dynamically according to the extent of peripheral nerve injury. This suggests that the activation of Dys and the deactivation of BF reflect the cortical representation of pain.

**Figure 3.**
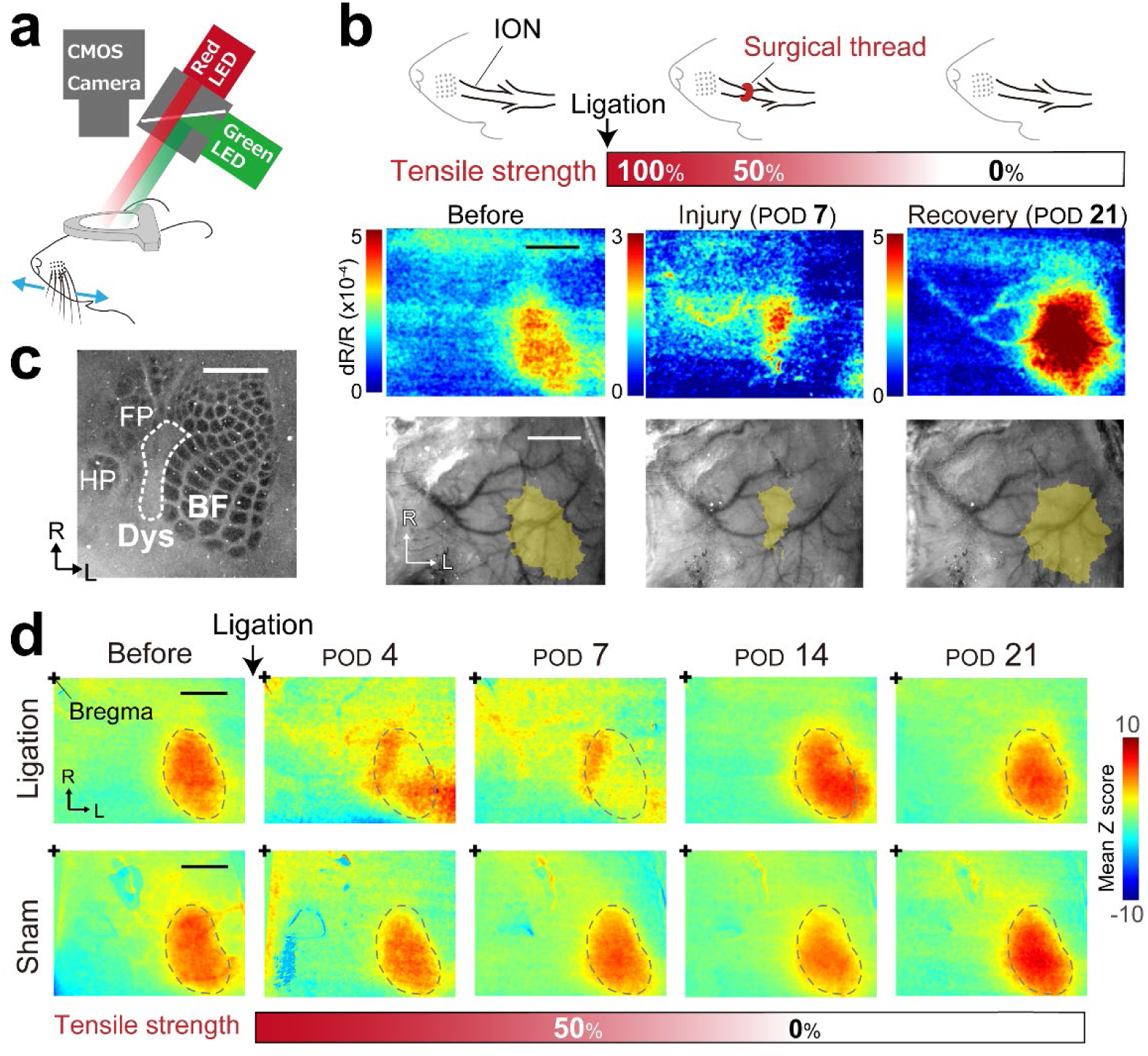
Tactile activated region was shifted from the barrel field to the dysgranular region during peripheral nerve injury. **a**, Schematic showing intrinsic signal imaging during whisker stimulation by the piezo device. Red LED was used for intrinsic signal imaging, and green LED was used for obtaining the vessel pattern of the brain surface. **b**, *Top*, Schematic of the time course for infraorbital nerve (ION) ligation by an absorbable surgical thread. *Middle*, Typical examples of the intrinsic signal images of an animal. *Bottom*, Overlaid images of the signal region (yellow) and vessel pattern used for alignment. **c**, Cytochrome c oxidase staining of a tangential section of S1 L4. FP, forepaw; HP, hindpaw. **d**, Averaged z-scored images (ligation group, *n* = 5; sham group, *n* = 3). The dotted areas indicate BF activated by whisker stimulation before ligation. R, rostral; L, lateral; POD, postoperative day. Scale bars, 1 mm.

### Dys is involved in generating pain behavior

Finally, we examined whether Dys is involved in pain behavior, such as escape from harmful stimuli. For this, we monitored the behaviors of head-restrained animals freely moving on a spherical treadmill^34^ in response to an innocuous or noxious infrared (IR) laser applied to the left whisker pad (Fig. 4a). Application of the IR laser for 500 and 1,500 ms increases the skin temperature to 39°C and 52°C, respectively^35^, which we thus refer to as innocuous heat (innH) and noxH, respectively. In response to noxH, the traveling speed increased until 5 s after the onset of the IR laser stimulus as the mice attempted to escape from noxious input (Fig. 4b–d and Extended Data Fig. 7a). The animals also exhibited eye blink and tightening (Extended Data Fig. 7a), which are considered expressions of pain^36–38^. Thus, this system was suitable for quantifying pain behaviors in response to noxH.

**Figure 4.**
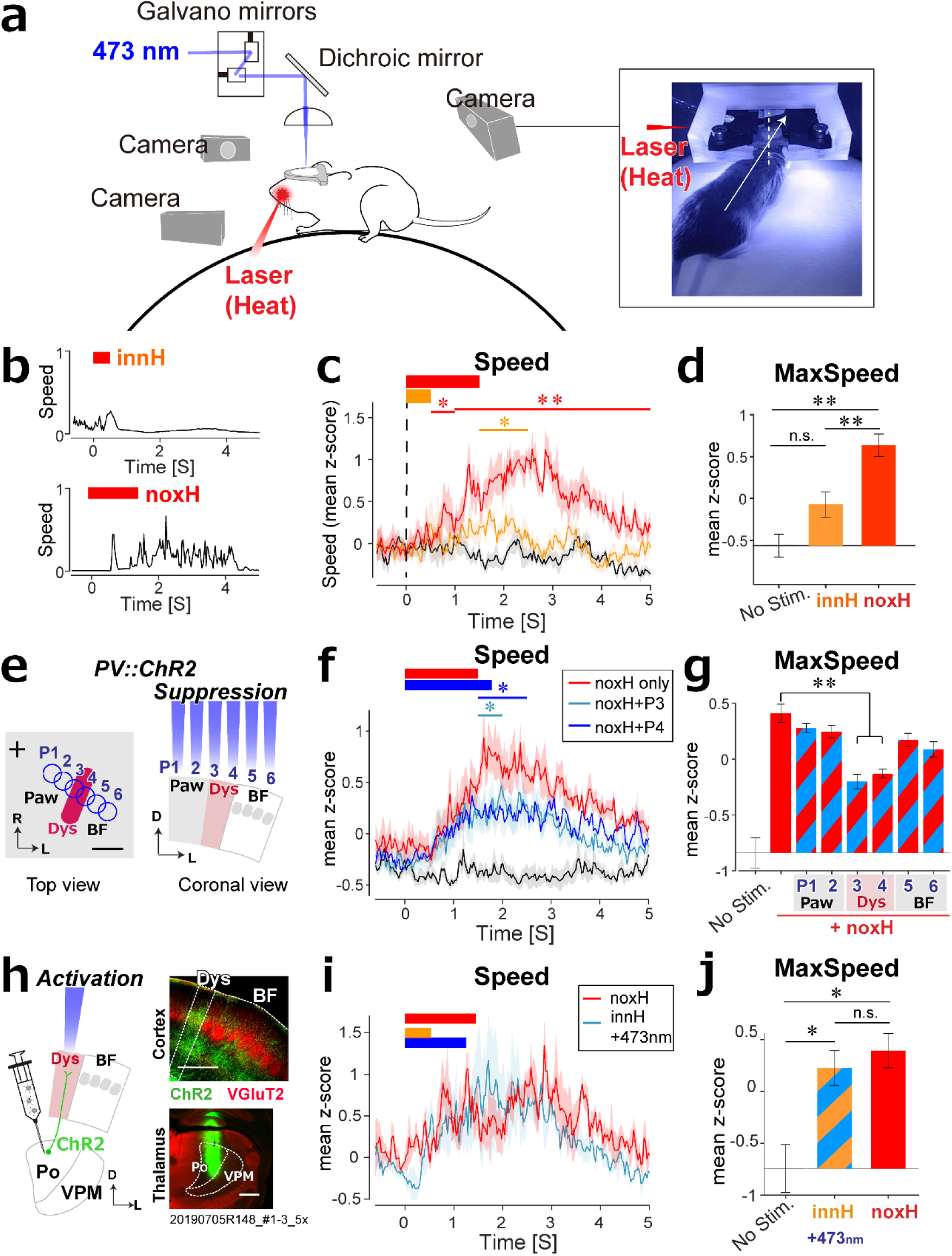
Dysgranular region activity drives nocifensive escape behavior. **a,** Schematic for monitoring escape behaviors induced by applying an 808 nm infrared (IR) laser to the whisker pad of a head-restrained freely moving animal on a spherical treadmill. Motion direction and the speed were monitored by a camera at the back, and facial expression and forelimb movement were monitored by side cameras. **b,** Examples of escape speed in response to 500 and 1,500 ms IR stimulation, corresponding to 0.09 (innocuous heat, innH) and 0.27 J/mm^2^ (noxH), respectively. **c**, Average speed profiles with (innH, orange; noxH, red) and without (black) IR stimulation (*n* = 6 mice). **d**, Maximum speeds induced by IR stimulation. **e,** Scheme for silencing six S1 positions (430 μm apart) via 473 nm photoactivation of channelrhodopsin-2 (ChR2) during noxH stimulation. Scale bar, 1 mm. +, bregma; R, rostral; L, lateral; D, dorsal. **f**, Average speed profiles show that optogenetic suppression at P3 (blue) and P4 (cyan) significantly reduced the escape speed to noxH (*n* = 6 mice). **g**, Mean z-scores of the maximum speed. **h**, Scheme for activating the thalamocortical fibers from the posterior nucleus (Po) and an example of ChR2 expression in Po and in the thalamocortical fibers in Dys. Scale bars, 500 μm. **i**, Average speed profile for optogenetic activation with innH matched that for noxH (*n* = 3 mice). **j**, Mean z-scores of the maximum speed. n.s., not significant; \**P* < 0.05, \*\**P* < 0.01, \*\*\**P* < 0.001; one-way ANOVA followed by Tukey–Kramer test. Shading and error bars indicate SEMs for mice. Osaki et al., (Supplementary table)

Using this system, we determined the effect of modulating Dys activity on pain behavior. We monitored pain behaviors induced by noxH during optogenetic suppression of various cortical areas, including Dys (Fig. 4e). For these experiments, we did not assess eye blink and tightening, which involve reflex actions via the brainstem or the cerebellum^39^. We used a transgenic line that expresses channelrhodopsin-2 (ChR2)-EYFP in parvalbumin (PV)-expressing interneurons (PV-Cre × Ai32), and photoactivated PV interneurons to inhibit cortical pyramidal neurons locally^40^(Fig. 4e). Photoinhibition of Dys significantly decreased the escape speed in response to noxH (for 1.5 to 2 s in position 3 (P3) and P4, *P* < 0.05; and for 2 to 2.5 s for P4, *n* = 6, Fig. 4f); photoinhibition of other S1 regions, such as BF or paw regions, had no effect (*P* > 0.05, Extended Data Fig. 8). Similar trends were observed for the maximal speed (*P* < 0.01 at P3 and P4; not significant at other positions, Fig. 4g), and for changes in the escape direction and distance (Extended Data Fig. 7b). In control mice that did not express ChR2, blue laser stimulation did not affect the escape speed (Extended Data Fig. 9). These results indicate that optogenetic local suppression of Dys neurons reduced noxH evoked pain behaviors.

Conversely, pain behaviors were also affected by Dys activation. Because the medial posterior nucleus (Po) neurons are thought to bring nociceptive information from the thalamus to S1^11^, we confirmed the connection from Po neurons to Dys by retro- and antero-grade tracers (Extended Data Fig. 10) ^16^ and utilised mice with virus-induced expression of ChR2 (AAV9-hSyn-ChR2(H134R)-EYFP) in Po neurons^16^. Photoactivation of Dys (Fig. 4h) in response to innH resulted in treadmill speed profiles resembling those in response to the noxH condition (Fig. 4i). Similarly, the maximal speed with innH condition paired with Dys photoactivation was not different from that in response to the noxH condition (*P* = 0.99, Fig. 4j). Photoactivation of Dys increased the escape speed at 2.5–3 s after innH onset compared with that without photoactivation (*P* < 0.001, *n* = 3, Extended Data Fig. 11c, d). Notably, this same increase was observed with ChR2 activation of Dys in the absence of innH (*P* = 0.016, Extended Data Fig. 11b). Similar trends were also observed in the escape direction and distance (Extended Data Fig. 7c). This suggests that ChR2-mediated photoactivation of Dys enhanced pain behaviors. In control mice that did not express ChR2 in Po, blue laser stimulation had no effect (Extended Data Fig. 12). In summary, the optogenetic studies demonstrate that Dys activation induces pain behaviors in response to innH, whereas Dys inhibition reduces them, revealing the role of Dys in generating pain behavior.

## Discussion

The results of this study reveal that Dys shows much higher selectivity to noxious heat input, whereas BF is more selective to tactile input. In particular, nociceptive information is processed separately from tactile information in Dys L2/3. Dys is also responsive to neuropathic pain. Reflecting the spatially-distinct representation of nociception, optogenetic suppression of Dys activity reduced noxious heat-evoked pain behaviors, whereas the same manipulation in BF showed no behavioral effect. These results indicate that Dys is mainly involved in nociception and in generating pain behaviors in S1.

### Separation and interaction of nociceptive and tactile processing

Previous studies have reported that nociresponsive cortical neurons are located in the deeper layers of BF^11,22,23^ and Dys^24^, but the functional differences between the two subregions remain unknown. Here, we demonstrate that Dys and BF have clearly segregated roles in nociceptive and tactile processing, especially in L2/3. Our neurophysiological data further show that noxH input suppresses neural activity in BF L2/3. This suppression may be the neural basis for disrupting the acuity of tactile sensation during pain^29,30^.

In contrast to that in L2/3, the segregation of nociceptive and tactile information processing was less clear in L5, with larger proportions of integrative neurons in both regions. However, the preference for tactile stimuli in L5 BF neurons was maintained; BF neurons showed greater selectivity to tactile input than Dys neurons (Extended Data Fig. 4c), while tactile input tended to suppress neural activity in Dys (Fig. 2f). Because L5 neurons are suggested to modulate nociceptive input through the corticofugal pathway^14,15^, BF L5 neurons might relate to touch-induced pain relief under normal conditions^31^, whereas Dys L5 neurons might contribute to mechanical allodynia (Fig. 3).

The notably finding in the present study is that Dys, but not BF, is involved in generating pain behavior (Fig.4). Several lines of anatomical evidence suggest that Dys closely relates to motor function. For instance, Dys receives proprioceptive inputs via Po^16,20,21^ and projects to the primary motor cortex^41^. Therefore, it is expected that somatic nociceptive information may integrate with proprioception in Dys. This combined information may exert influence over the motor pathway, which leads to proper escape behavior from noxious inputs. Because the descending projections from Dys terminate in areas distinguishable from those from S1 cutaneous areas^42^, the functional differences of the outputs from Dys and BF should be examined in more detail in future studies.

### Possibility of S1 intervention for pain relief based on S1 functional structure

Although deep brain stimulation of the somatosensory thalamus, the ventral posterolateral nucleus and ventral posteromedial nucleus, had been utilized to treat chronic pain^43^, it is not currently used as a medical treatment because of its limited therapeutic effect^44,45^. The reason why the limitation may be the fact that the sensory pathways are not selectively stimulated. Therefore, we guess that the selective inhibition of nociresponsive region in S1 could overcome it and have more therapeutic effects.

Neurons in primate area 3a, which is also a dysgranular region in cytoarchitecture^46^, responds to noxious ^46,47^ and proprioceptive inputs^48^. Thus, primate area 3a could be an evolutionally homologue to rodent Dys. Considering Dys inhibition reduced pain behavior (Fig. 4), inhibition of area 3a might work as pain relief. However, area 3a is buried in the fundus of the central sulcus in humans^49^, the selective intervention of area 3a is challenging and needs to be resolved for effective treatment of chronic pain in the future^46,47^. Nevertheless, our results provide evidence that the distinctive nociceptive region in S1 is a potential therapeutic target for pain relief.

## Acknowledgments

We thank Sachie Sekino and Yumi Tani for their excellent technical assistance. This research was supported by JSPS KAKENHI grant numbers JP15K21387, JP15H01667, JP17H05912, JP18K14854, and The Uehara Memorial Foundation to H.O. This research was also partially supported by JSPS KAKENHI grant numbers JP20H05481, JP20H05916, JP20K21508, JP17H05752 and the SHISEIKAI Scholarship Fund for Basic Researcher of Medical Science, Keiko Watanabe Award to M.M. and the program for Brain Mapping by Integrated Neurotechnologies for Disease Studies (Brain/MINDS) from the Japan Agency for Medical Research and Development, AMED, under grant number JP19dm0207057.

## Author contributions

HO and MM designed the experiments. HO, MK, and YU performed the experiments and analyzed the data. HO and MM wrote the original draft.

## Methods

### Animals and surgery

All surgical procedures and postoperative care were performed according to the guidelines of the Animal Care and Use Committee of Tokyo Women’s Medical University. The animal experiment was approved under number AE19-109. C57BL/6N (Sankyo Lab. Service Corp., Tokyo, Japan), PV-Cre (JAX stock #008069), and Ai32 (Rosa-CAG-LSL-ChR2[H134R]-EYFP-WPRE; JAX stock #012569) mouse lines were used in this study. PV-Cre mice were crossed with Ai32 mice, and the resulting mouse line was designated PV-ChR2. Male mice of 8 weeks or older in age were used. The animals were group-housed in a cage maintained at 23°C ± 1°C with a 12 h light/dark cycle, and all behavioral tests were performed during the dark period. Every effort was made to minimize the number of mice used and their suffering in this study. For surgical procedures, each animal was anesthetized with an intraperitoneal injection of ketamine (100 mg/kg body weight) and xylazine (16 mg/kg body weight), and isoflurane was supplemented to maintain the anesthesia. Lidocaine was applied subcutaneously at the incision site and to the wound margins for topical anesthesia. For intrinsic signal optical imaging, electrophysiological recordings, and behavioral testing on a spherical treadmill, a custom-built headplate was attached to the skull with dental acrylic clear resin (Super-Bond; Sun Medical, Shiga, Japan). The head plate-implanted animals were returned to the home cage and allowed to recover from the surgery for at least 4 days.

### Capsaicin injection

For capsaicin injections into the whisker pad, mice were anaesthetized with 2% isoflurane. Capsaicin (Wako Pure Chemical Industries, Ltd., Osaka, Japan) was dissolved in 100% ethanol (229 ul), and then mixed with 7% Tween 80 in saline (3.04 ml). Capsaicin (10 mM, 50 μl) was injected into the left whisker pad. A solution containing 100% ethanol, 7% Tween 80, and saline was used as the vehicle^1^. One hour after the injection, the animals were perfused with 4% paraformaldehyde with picric acid followed by c-Fos immunohistochemistry using the diaminobenzidine (DAB) protocol described below.

### c-Fos immunohistochemistry using DAB

After the brains were postfixed, 50-μm-thick slices were made, and alternate slices were reacted with cytochrome c oxidase to identify BF, and with c-Fos immunohistochemistry. For c-Fos immunohistochemistry, the slices were processed with 1% H_2_O_2_ in phosphate buffer to deactivate the intrinsic peroxidase and then incubated with an anti-c-Fos antibody (1:10,000, rabbit; Merck KGaA, Darmstadt, Germany) in 10% normal goat serum in phosphate-buffered saline with 0.3% Triton X-100 (PBS-X) at 4°C overnight. The slices were then incubated with a biotinylated goat anti-rabbit IgG antibody (1:200; Vector Laboratories, Burlingame, CA, USA) and reacted with avidin-biotin-peroxidase complex (ABC kit; Vector Laboratories). The slices were incubated in a DAB solution (0.02% DAB, 0.3% nickel ammonium sulfate in Tris-buffered saline) and 1% H_2_O_2_ for visualization. Images were acquired with an upright microscope with a charge-coupled-device camera (DP70; Olympus, Tokyo, Japan). The neurons expressing c-Fos were counted by a custom-written MATLAB (MathWorks, Natick, MA, USA) program with edge detection by a Sobel filter followed by binarisation. The numbers of c-Fos-positive neurons in two different slices were averaged to minimize selection bias.

### Electrophysiological recording

For electrophysiological recordings from Dys and BF regions, each mouse was anaesthetized with isoflurane (0.4–0.8%) supplemented with an intraperitoneal injection of chlorprothixene hydrochloride (2 mg/kg body weight)^2^. As the nociceptive response depends on the level of anaesthesia^3^, the respiration rate (70–120 cycles/s) was monitored to control the level of anesthesia.

After identifying the border between Dys and BF regions according to intrinsic signal imaging, 32-channel four-shank electrodes (A4x8 or Buzsaki32; NeuroNexus, Ann Arbor, MI, USA) were inserted into S1 to record from Dys and BF regions simultaneously. Raw electrical signals were amplified and digitized at 40 kHz (Plexon, Dallas, TX, USA) and then processed for spike sorting. The spike sorting comprised automated spike detection and clustering using Klusta followed by manual sorting using Kwik GUI^4^. First, noise artifacts determined from the waveform were extracted. Second, multiunit activity (MUA) was determined from the waveform with low amplitude and without a refractory period (>2 ms) in autocorrelograms. Third, after merging and/or splitting clusters using auto- and cross-correlograms and principal-component features, single-unit activity (SUA), which has a clear refractory period (>2 ms) in autocorrelograms, or MUA was determined. To estimate the depth of recorded neurons, the maximum amplitudes of waveforms from each probe were compared and determined for the nearest probe for each SUA/MUA. For MUA analysis, SUA is included in MUA (*n* = 8 animals, 128 probe sites for simultaneously recorded MUA analysis).

### Calculation of the S/N for noxious and tactile stimuli

We calculated the S/N to identify the area or cell properties and assessed selectivity to noxious heat and tactile stimuli. For the S/Ns for the noxious heat response, the response for 500 ms over 45℃ of the Peltier device (45–50℃, beige shading, Fig. 1e) was counted as the S response. The innocuous heat response (for 1,000 ms until 45℃ corresponding to 33–45℃, grey shading, Fig. 1e) was counted as the N response. This definition helped to select noxious heat-selective neurons by excluding the temperature-coding neurons.

For tactile stimulus, the response for 30 ms after the onset of the whisker deflection (blue arrows, Fig. 2a, right; Extended Data Fig. 3) was counted as the S response. The response for 60 ms until the onset of the whisker deflection was counted as the N response (Extended Data Fig. 3). All deflections were used for this calculation (Extended Data Fig. 3). This definition helped to detect how the neuron responds to sequential whisker deflection^5^.

### Formalin injection and c-Fos immunofluorescence

Each mouse received an intraplantar injection into the left hind paw of 50 μl of a 5.0% formalin solution using a 27-gauge needle. One hour later, the animals were perfused with 4% paraformaldehyde with picric acid. The brains were removed, and the right hemispheres were flattened between two glass slides. After the brains were postfixed, tangential 50-μm-thick sections were made, followed by c-Fos and vesicular glutamate transporter type 2 (vGluT2) immunostaining and Nissl staining. Briefly, the sections were incubated with anti-c-Fos antibody (1:5,000, rabbit; Merck KGaA) and anti-vGluT2 antibody (1:500, vGluT2-GP-Af810; Frontier Science, Ishikari, Japan) in 10% normal donkey serum in PBS-X at 4℃ overnight. The sections were then incubated with the following secondary antibodies: anti-rabbit antibody (Alexa Fluor 568 conjugated, 1:1,000, donkey; Invitrogen) and anti-guinea pig antibody (Alexa Fluor 647 conjugated, 1:200, donkey; Jackson ImmunoResearch, West Grove, PA, USA). Images were acquired with an epifluorescence microscope (Axio Scope.A1; Carl Zeiss) with a cooled charge-coupled-device camera (RS 6.1; Quantum Scientific Imaging) and by using μManager (http://www.micro-manager.org) and ImageJ software (https://imagej.nih.gov/ij). The neurons expressing c-Fos were counted by a custom-written MATLAB program with edge detection by a Sobel filter, followed by binarisation.

### Anterograde/retrograde labelling

For retrograde labeling, cholera toxin subunit B conjugated with Alexa Fluor 555 (0.2%) or with Alexa Fluor 488 (1%) were applied after intrinsic signal imaging to identify the Dys or BF region, respectively. Three days later, the mouse was deeply anaesthetized with sodium pentobarbital (60 mg/kg body weight, intraperitoneally) and transcardially perfused with a fixative solution (4% paraformaldehyde and 0.2% picric acid in 0.1 M phosphate buffer). The brains were removed and cut coronally into 50 μm sections. Sections were incubated overnight with a guinea pig polyclonal antibody against vGluT2 (1:500; vGluT2-GP-Af810) followed by NeuroTrace 435/455 (1:100; Thermo Fisher Scientific, Waltham, MA, USA).

For anterograde labeling, biotinylated dextran amine (molecular weight: 10,000, 10% in saline; Thermo Fisher Scientific) was injected into the Po (1.7 mm posterior to bregma and 1.3 mm lateral to the midline). Seven days later, the mouse was deeply anesthetized and brain sections were cut, as described above. Sections were incubated overnight with vGluT2-GP-Af810 antibody followed by Alexa Fluor 594-conjugated secondary antibody (1:500; Jackson ImmunoResearch), Alexa Fluor 488-conjugated streptavidin (Thermo Fisher Scientific), and subsequently with NeuroTrace 435/455 (1:100; Thermo Fisher Scientific).

### Intrinsic signal optical imaging

At least 4 days after the head plate implantation, intrinsic signal imaging was performed to measure the responses from BF. The mouse was anesthetized as described above in “Electrophysiological recording”. The respiration rate and heart rate were monitored via a video-based respiration monitor with a 30 Hz web camera (C920; Logitech, Lausanne, Switzerland) and an acceleration monitor on the back of the animal. Intrinsic signal images were obtained with a CMOS camera (MV1-D1024E-160-CL; Photonfocus, Lachen, Switzerland) with the tandem lens of an achromatic doublet (Thorlabs, Newton, NJ, USA) and long-pass and short-pass filters (BLP01-488R-25 and FF01-650/SP-25; Semrock, Rochester, NY, USA). Frames were acquired at a rate of 20 Hz, and the frame size was 600 × 500 pixels and represented 5.5 × 4.5 mm of cortical area. The brain surface was illuminated by a green LED (M530L3; Thorlabs) to obtain the vessel pattern or a red LED (M617L3; Thorlabs) to obtain the intrinsic signal. Images were recorded through the skull covered with dental acrylic clear resin. The dental acrylic resin was covered with a nail topcoat and silicone immersion oil (Olympus) to reduce glare.

Whisker stimulations as tactile stimuli were generated using a piezoelectric device as described previously^5^. To visualize the cortical response to tactile stimuli, we calculated the reflectance ratio in each frame (dR/R, where dR is the difference of reflectance R from the base image that is the average from 20 frames taken before stimulus onset). To map the change in cortical activity, images taken on different experimental days were aligned according to vessel patterns using a custom-written MATLAB program. For population analysis, we calculated the z-scored image from dR/R using the following equation:

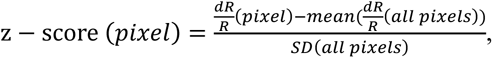

where SD is the standard deviation. The positions of bregma were used for alignment between the animals.

### ION ligation and behavioral assay using von Frey filaments

To produce a neuropathic pain model, the ION was tightly ligated by a surgical thread (Vicryl Rapide; Ethicon, Bridgewater, NJ, USA). This thread enables us to observe changes in BF activity during both nerve injury and recovery, as the tensile strength of the thread gradually reduces inside the body (within 2–3 weeks); the tensile strength is reduced to almost 50% at POD 7 and 0% at POD 21, corresponding to the phases of nerve injury and recovery, respectively. For the behavioral assay of neuropathic pain, the animals were trained to enter a 50 ml tube with a custom-made tube holder. Behavioral training began after the mice had restricted access to water (1 ml/day) before at least 7 days of left ION ligation or sham operation. The water was restricted to 1.5 ml for 1 day during the behavioral experiment. The animals were trained to enter the tube and keep their snouts protruded through a hole to drink water. While the animals were drinking, the left whisker pad was stimulated by von Frey filaments (1.4, 2, 4, 6, 8, and 10 g; Ugo Basile, Varese, Italy) to measure the escape threshold^6^. During stimulation by the filament, visual information was blocked by a black cover (Extended Data Fig. 6).

### Environmental enrichment

At 7 or 21 days after ION ligation and 7 days after sham operation, the mice were placed into an enriched environment for 1 h to enhance whisking while exploring several objects (Extended Data Fig. 6). Then, the mice were perfused with 4% paraformaldehyde with picric acid followed by c-Fos immunohistochemistry using the DAB protocol described above.

### Measurement and analysis for pain behavior on a spherical treadmill

Mice were first trained to enter a tube to obtain a water reward for 3–4 days. Mice were then head-fixed and free to run on a spherical treadmill (Ø 30 cm) under a white noise sound condition^7^ for 3–4 days. After this, the behaviors of the mice towards the IR laser stimulus on the left whisker pad were monitored. For the noxious heat stimulus, an IR diode laser (Ø 1 mm on the whisker pad, wavelength of 808 nm, SSL-808-1000-10TM-D-LED; Shanghai Sanctity Laser Technology Co., Ltd., Shanghai, China) was used. The stimulus duration was set to 500 or 1,500 ms, corresponding to 0.09 or 0.27 J/mm^2^, respectively, to increase the skin temperature to 39°C or 52°C^8^. At the start and end of each session, the animals obtained a water reward on the treadmill but not during the stimulus sessions. Each session was composed of various stimulus conditions: each condition was randomly chosen and presented to the animal five times in one session. The animals were imaged with three cameras at 30 Hz (the IR filter on the CMOS sensor was removed, C922; Logitech, Lausanne, Switzerland) set behind and to each side of the animal to record behaviors such as escape direction, speed, moving distance, eyeblink, and left forelimb movement. These parameters were analyzed by calculating the difference of the region of interest of each parameter frame by frame. These parameters were calculated into z-scores for comparisons among the animals by a custom-written MATLAB code. If the animal ran continuously on the treadmill and did not show any difference in the maximum speed between trials, the session was excluded from the analysis. At least 2 sessions were used for calculation of z-scores. A notch filter (808 nm OD4 notch filter, 86-702; Edmund Optics) was placed in front of the left side camera to prevent sensor white-out and to identify the precise position and size of the IR laser stimulation. Custom-made LED illuminators (940 nm) were placed in front of each camera to illuminate the animals.

### Optogenetic activation and suppression of Dys

To activate the thalamocortical fibers from Po into Dys, a virus (AAV9-hSyn-hChR2(H134R)-EYFP) was injected into the right Po (1.7–1.9 mm caudal and 1.2 mm lateral to bregma; depth, 2,800 and 3,000 μm from the brain surface; 50 nl at each depth) (QSI; Stoelting, Wood Dale, IL, USA). A blue laser (473 nm; Lasos, Germany) was coupled to an optic fiber cable (Ø 200 μm; Thorlabs). The output of the optic fiber and the surface of the cortex were placed on conjugate planes using a fiber port and an achromatic lens. The *x* and *y* Galvano mirrors (Galvano scanning system, GVS002; Thorlabs) were placed in the infinity space to control the position of the stimulus. A dichroic mirror was also placed in the infinity space, and the reflected light was focused onto the sensor of a CMOS camera (Grasshopper3; FLIR). This design enabled monitoring of the precise location of the stimulated site^9^.

An optical chopper (pulse width, 2.5 ms, MC2000B; Thorlabs) was used to activate PV interneurons with a 40 Hz pulse^10^. To investigate the suppressive effect of Dys activity, the six positions covering Dys were stimulated by a blue laser focused on the brain surface by an achromatic doublet lens (AC254-60-A; Thorlabs). The centers of the stimulation sites were 430 µm apart, the spot diameter was 1.5 mm, and the power was ∼0.9 mW per location.

### Code and data availability

All codes used for analysis in this study were implemented in MATLAB. The codes and the data are available from the corresponding author upon reasonable request.

---

**Supplementary Table 1.** Capsaicin injection into the whisker pad increases the number of c-Fos-positive neurons in Dys L4.

| Layer | Dys |  |  | BF |  |  |
| --- | --- | --- | --- | --- | --- | --- |
|  | Density of c-Fos<sup>+</sup> neurons (cells/10<sup>4</sup> μm<sup>2</sup>) |  | *P* value | Density of c-Fos<sup>+</sup> neurons (cells/10<sup>4</sup> μm<sup>2</sup>) |  | *P* value |
|  | Capsaicin | Vehicle |  | Capsaicin | Vehicle |  |
| L23 | 10.5±2.3 | 8.5±0.6 | 0.99 | 13.1±1.8 | 13.0±0.4 | 1.0 |
| L4 | 26.4±2.5 | 11.8±2.0 | 1.2×10<sup>-5</sup> | 27.3±5.0 | 23.1±3.6 | 0.97 |
| L5a | 11.0±1.5 | 5.4±0.4 | 0.25 | 10.9±2.3 | 10.8±1.9 | 1.0 |
| L5b | 10.6±0.8 | 8.4±1.0 | 0.99 | 11.3±2.4 | 13.5±1.7 | 1.0 |
| L6 | 13.9±0.8 | 7.3±1.7 | 0.10 | 14.2±1.9 | 11.3±2.4 | 1.0 |

Values are means ± SEMs; *P* values are from Tukey–Kramer test (capsaicin, *n* = 3; vehicle, *n* = 4). The same data were used in the bar graphs in Fig. 1b.

**Extended Data Fig. 1.**
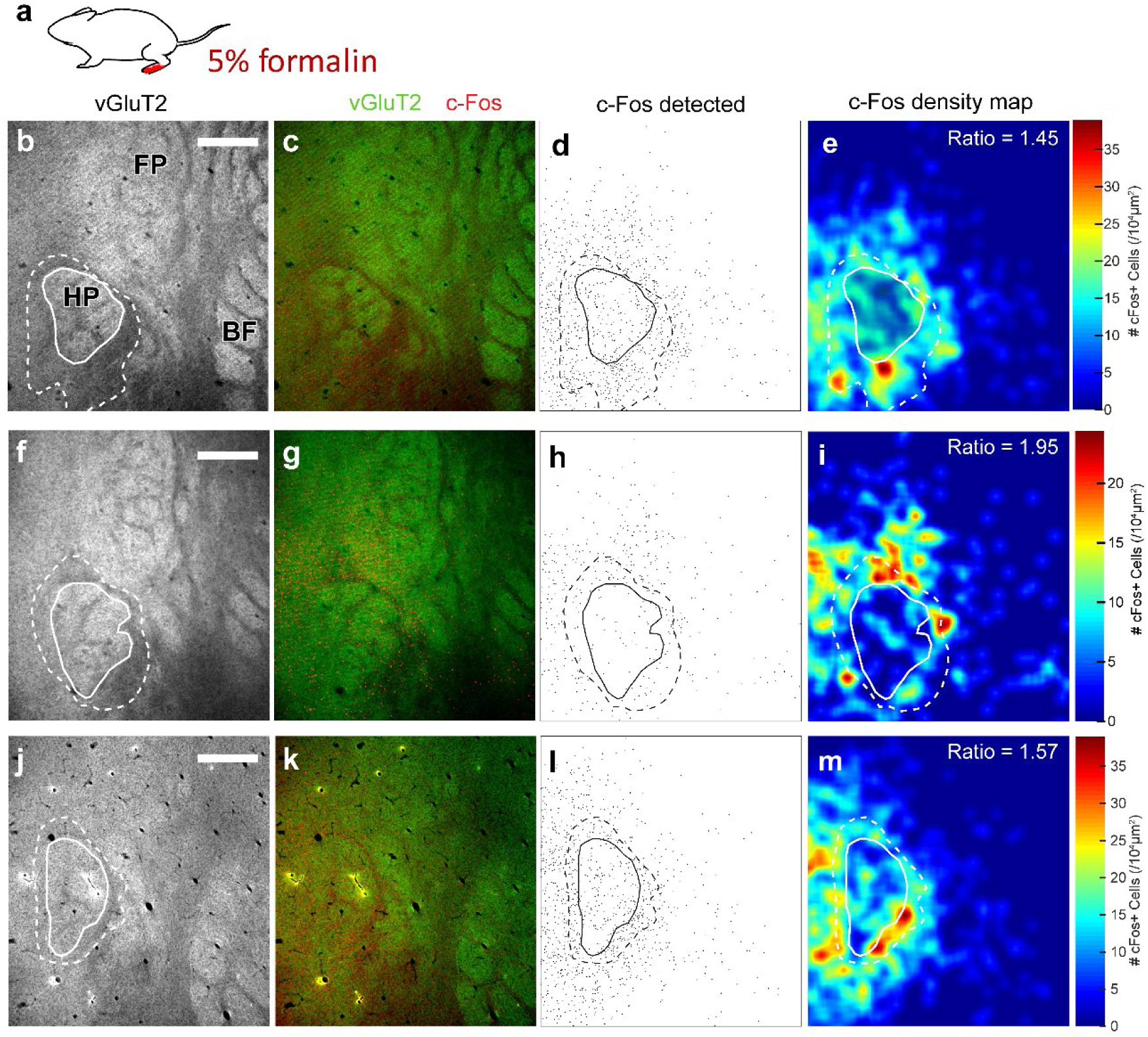
Noxious input into hind paw activates Dys L4 of the hindpaw area. **a**, Formalin (5%) was injected into the left hind paw. **b**,**f**,**j**, Tangential sections of S1 L4 from different mice. The granular areas of hindpaw (HP, white trace) and Dys (white dotted trace) were determined by immunostaining of the vesicular glutamate transporter-2 (vGluT2). FP, forepaw; BF, whisker barrel field. Scale bars, 400 μm. **c**,**g**,**k**, Co-immunostaining of vGluT2 (green) and c-Fos (red) after formalin injection into HP. **d**,**h**,**l**, Maps of c-Fos-positive neurons. **e**,**i**,**m**, Colour maps of c-Fos density overlayed with granular and dysgranular regions for the hind paws. Ratio of c-Fos-positive neurons (Dys/HP) of each mouse is shown.

**Extended Data Fig. 2.**
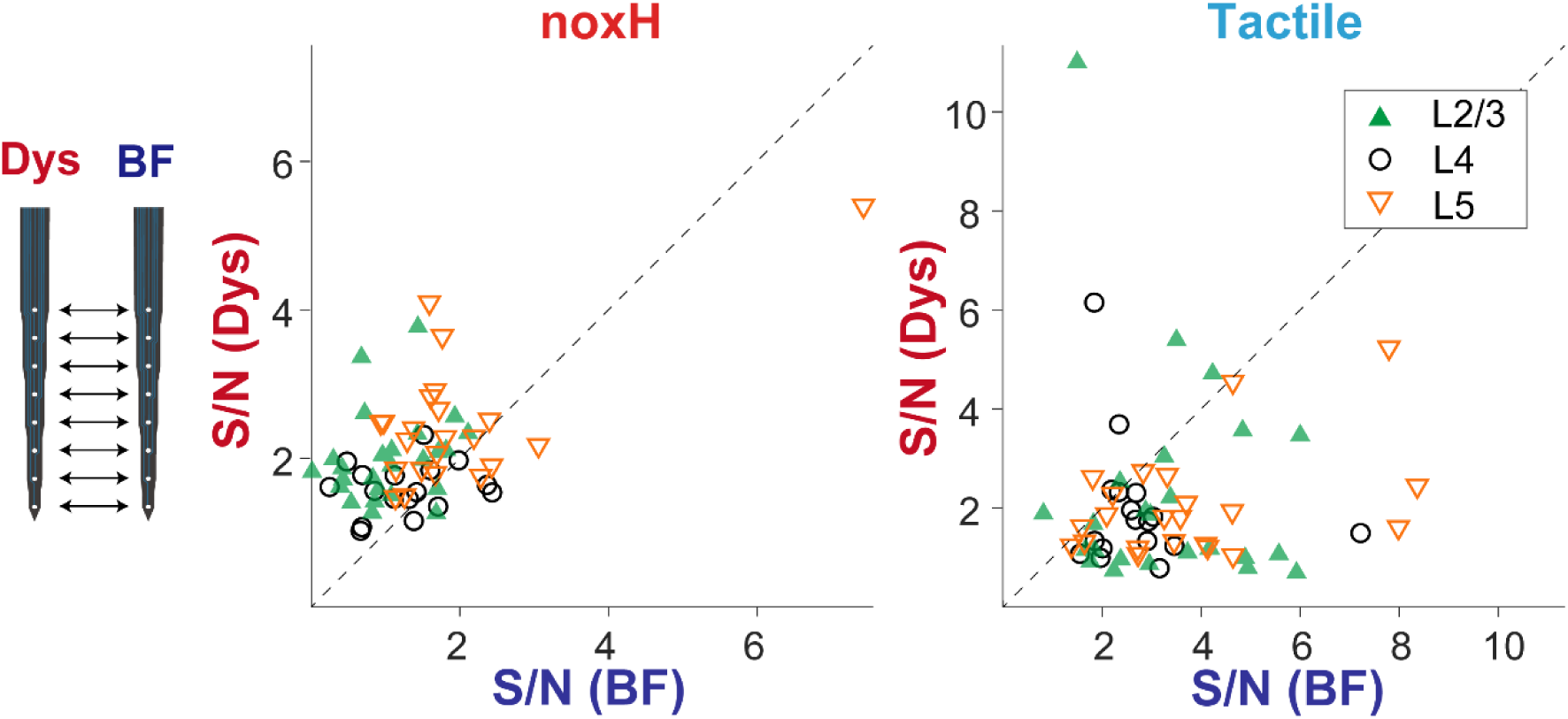
Simultaneously recorded multiunit activities show the difference of response properties between Dys and BF to noxH and tactile stimuli. The diagonal indicates unity. *Left*, Scatter plot of S/N to noxious heat (noxH) stimulus. *P* = 0.000023 for L2/3 (*n* = 25), 0.0056 for L4 (*n* = 17), and 0.062 for L5 (*n* = 22) by Wilcoxon signed-rank test. *Right*, Scatter plot of S/N to tactile stimulus. *P* = 0.00382 for L2/3 (*n* = 25), 0.028 for L4 (*n* = 17), and 0.00020 for L5 (*n* = 22) by Wilcoxon signed-rank test.

**Extended Data Fig. 3.**
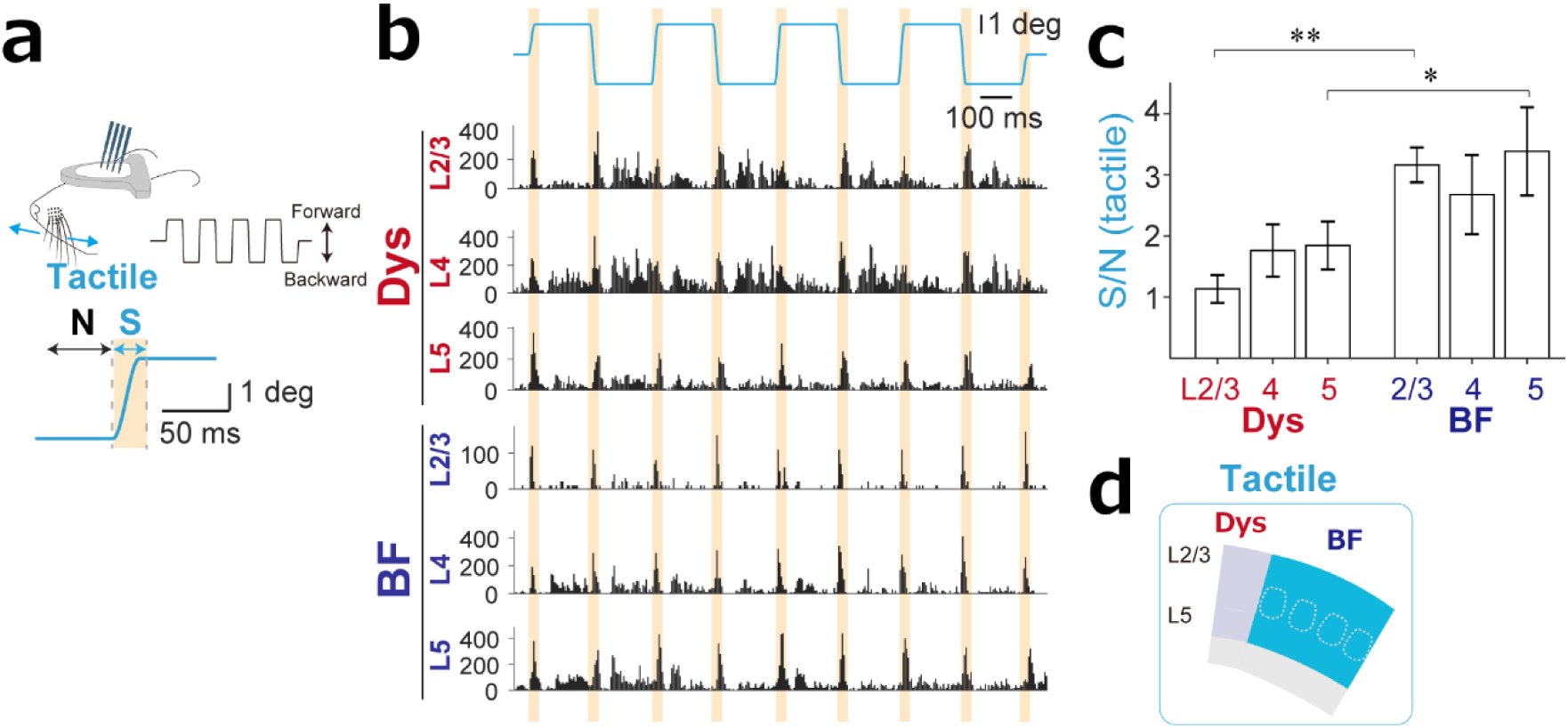
BF responds more selectively to tactile input than Dys. **a**, *Top*, Setup for recording responses to tactile stimulation of whiskers. The four-shank electrodes were inserted into Dys and BF. *Bottom*, Shaded area indicates the time used as signal (S) and noise (N) in S/N calculation of tactile input. **b,** Examples of perstimulus time histograms of multiunit activity to tactile stimulus in L2/3, 4, and 5 of Dys and BF recorded at the same time (see also Fig. 1e). The S/Ns to tactile stimuli were 1.1 (L2/3), 1.8 (L4), and 1.8 (L5) in Dys and 3.2 (L2/3), 2.7 (L4), and 3.4 (L5) in BF. **c,** S/Ns to a tactile stimulus were higher in BF in L2/3 (*P* = 5.4×10^-4^) and L5 (*P* = 0.031) (compare with Fig. 1e). \**P* < 0.05; \*\**P* < 0.01 by Kruskal–Wallis test followed by Dunn’s test; *n* = 8 animals. **d**, Summary diagram indicates that S/N for tactile stimuli was higher in BF than Dys.

**Extended Data Fig. 4.**
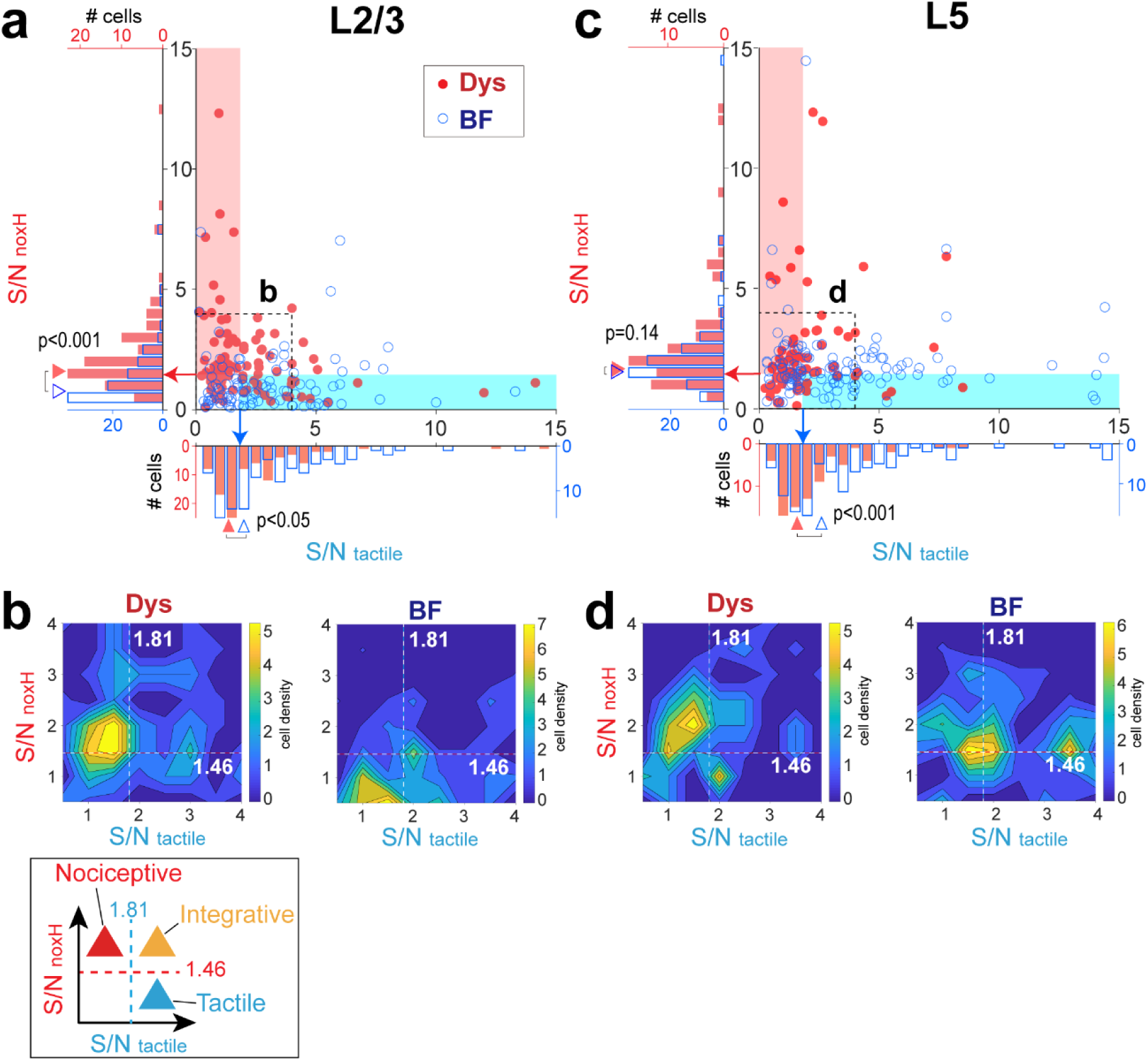
Segregation and integration of noxious heat and tactile information were observed in the scatter plots of S/Ns of all neurons. **a**, Scatter plot of S/Ns to a tactile stimulus against S/Ns to noxious heat stimulus (noxH) for L2/3 neurons. The neurons in the blue shaded area are the tactile input-preferring neurons, for which S/Ns to a tactile stimulus were higher than 1.81 (the median of all neurons, blue arrow), and the S/Ns to noxH were lower than 1.46 (the median of all neurons, red arrow). The neurons in the red shaded area are the noxious input-preferring neurons, for which S/Ns to a tactile stimulus were lower than 1.81, and the S/Ns to noxH were higher than 1.46. The distribution of S/Ns for noxH of Dys is significantly higher than that of BF (*P* = 4.82×10^-9^, two-sample Kolmogorov–Smirnov test). Red arrowhead on the *y* axis indicates the median S/N for noxH of Dys neurons, 1.60; blue arrowhead indicates median of BF neurons, 0.77. The distribution of S/Ns for tactile stimulus in Dys is significantly lower than that in BF (*P* = 0.012, two-sample Kolmogorov–Smirnov test). Red arrowhead on the *x* axis indicates the median S/N for tactile stimulation of Dys neurons, 1.36; blue arrowhead indicates median of BF neurons, 2.00. **b**, Density map of S/Ns to tactile stimulus against S/Ns to noxH of L2/3 neurons, with each S/N <4. **c**, Same as for panel b but for L5 neurons. The distributions are not significantly different from each other for S/Ns for noxH (*P* = 0.14, two-sample Kolmogorov–Smirnov test). Red arrowhead on the *y* axis indicates the median S/N for noxH of Dys neurons, 1.65; blue arrowhead indicates the median of BF neurons, 1.52. The distributions of S/Ns for tactile stimulation are different from each other (*P* = 0.00056, two-sample Kolmogorov–Smirnov test). The red arrowhead on the *x* axis indicates the median S/N for tactile stimulation of Dys neurons, 1.58; blue arrowhead indicates median of BF neurons, 2.57. **d**, Same as for panel c but for L5 neurons. *Bottom*, Cell classification (see also Fig. 2c).

**Extended Data Fig. 5.**
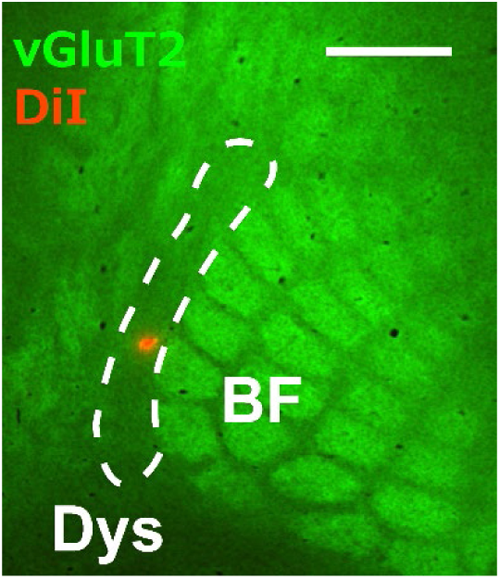
Region adjacent to BF was identified as Dys. An electrode stained with DiI (red) was inserted into the region adjacent to BF after intrinsic signal imaging was performed. Immunohistochemistry was performed on a tangential section of L4 to identify BF via vGluT2 (green). Scale bar, 500 μm.

**Extended Data Fig. 6.**
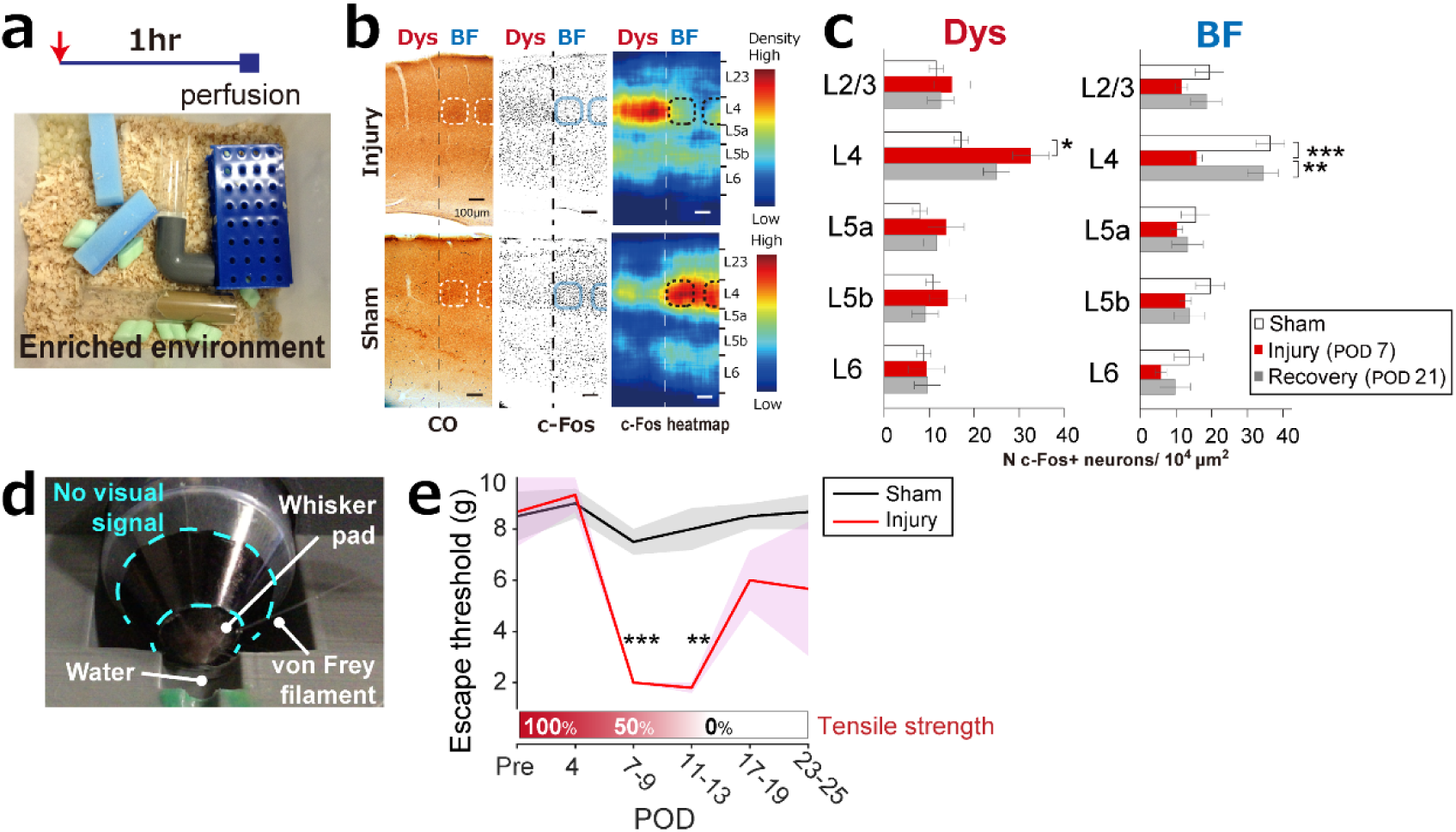
Infraorbital nerve ligation by absorbable surgical thread induced allodynia and activated Dys. **a-c**, Dysgranular area was activated under enriched environmental conditions during infraorbital nerve ligation. **a**, Animals were placed in the enriched environment for 1 h before perfusion. **b**, Examples of CO staining, c-Fos expression, and c-Fos density heat map 7 days after ligation/sham operation (POD7). **c**, L4 of Dys was activated at day 7. c-Fos expression in L4 of BF decreased at day 7 and recovered to sham level at day 21. \**P* = 0.0485, \*\**P* = 0.00318, \*\*\**P* = 0.000242 by one-way ANOVA followed by Tukey–Kramer test; sham, *n* = 4; injury, *n* = 3; recovery, *n* = 3. **d**, Setup for von Frey test. **e**, Mechanical allodynia was observed at 7–9 (\*\*\**P* = 0.0002) and 11–13 (\*\**P* = 0.0015) days after nerve ligation (one-way ANOVAs followed by Dunnett’s test). Error bars and shading indicate SEMs. Sham, *n* = 3; injury, *n* = 3.

**Extended Data Fig. 7.**
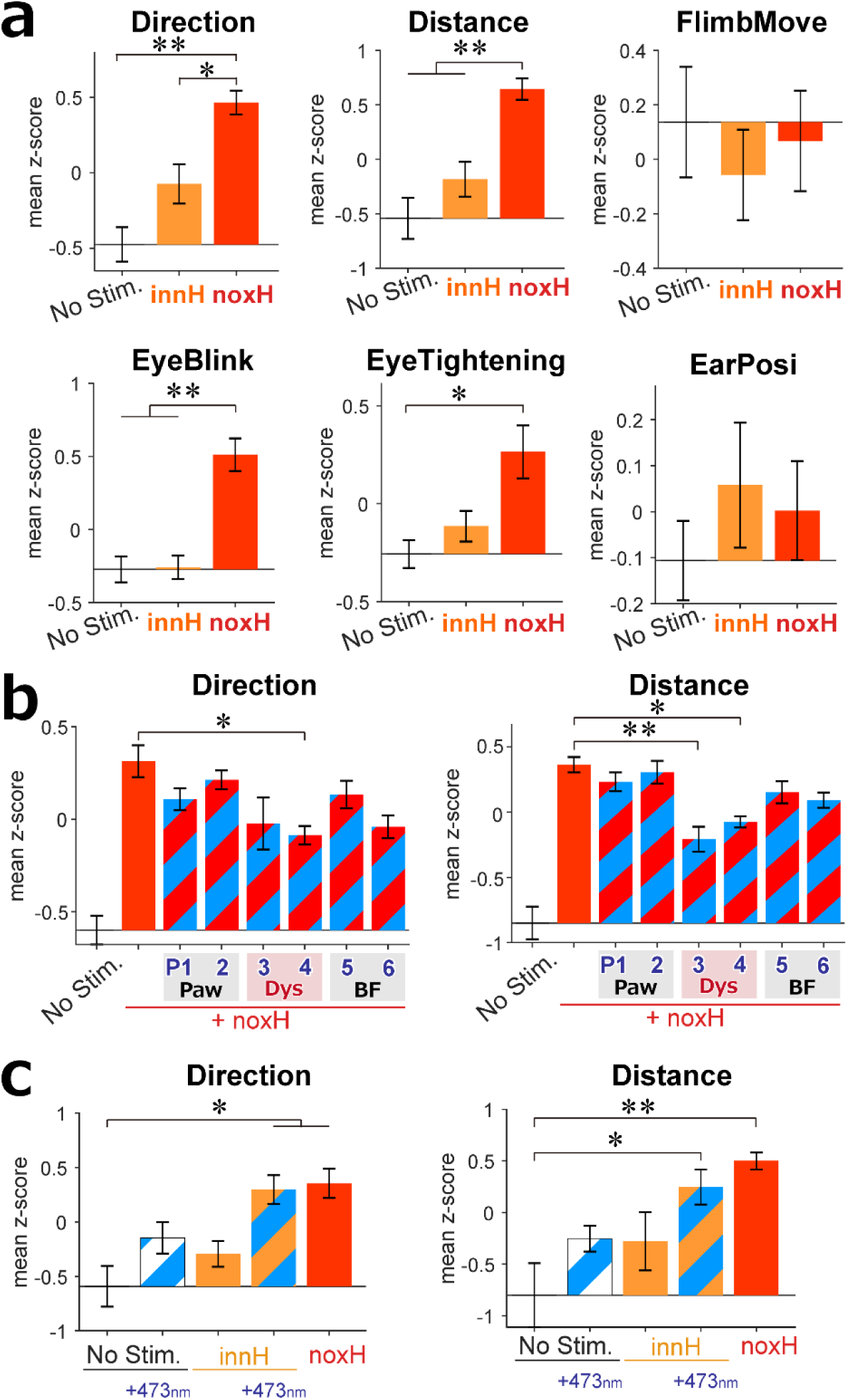
Nocifensive responses taken from the behavioural experiment using a spherical treadmill. **a**, Indices to measure nocifensive responses to IR laser exposure for 0 (no stim), 500 (innH), and 1,500 ms (noxH). Direction, the moving direction; a high z-score indicates the animal moved contralateral to the direction of the IR laser. Distance, the total distance traveled during 4 s after the onset of IR laser. FlimbMove, the number of touches or covering of the left whisker pad by the left forelimb. EyeBlink, the number of eye blinks. Eye tightening, sustained closure of left eyelid. Error bars indicate SEMs (*n* = 6). **b**, Study of the activation of PV interneurons in different anatomical positions (*n* = 6). **c**, Indices of the activation of thalamocortical fiber from Po after IR laser stimulation (*n* = 3). \**P* < 0.05, \*\**P* < 0.01 by one-way ANOVA followed by Tukey–Kramer test.

**Extended Data Fig. 8.**
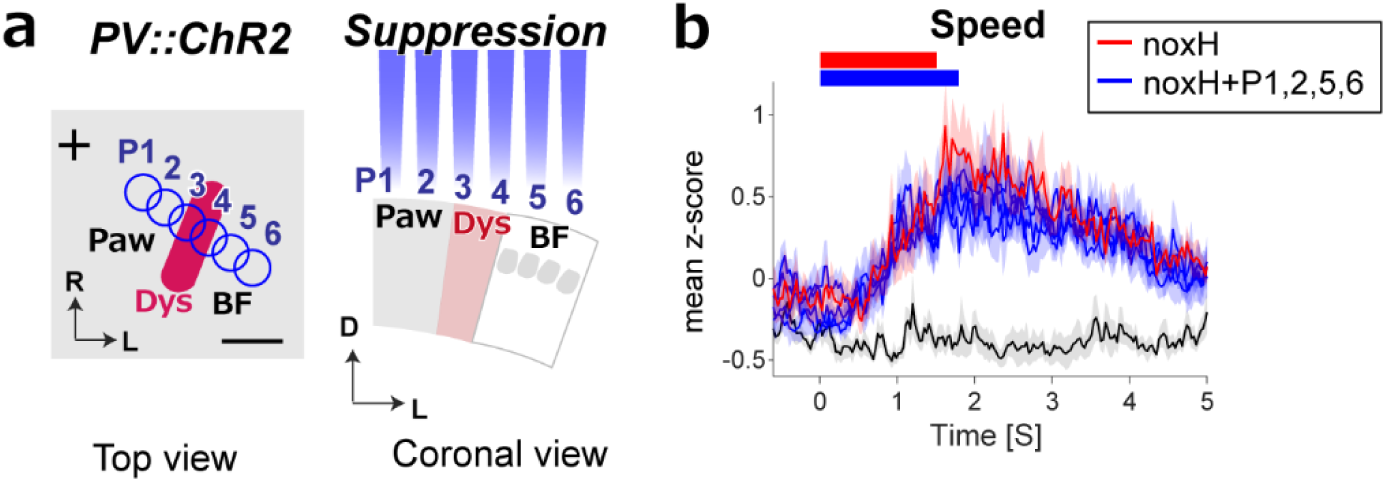
Blue light stimulation of BF did not reduce escape speed. **a**, Same setup as shown in Fig. 4e. **b**, Blue light stimulation of P1, 2, 5, and 6 did not reduce the escape speed to noxH. The difference between the IR-only condition and P1, 2, 5, 6 activation combined with IR stimulation was statistically insignificant at all time points (one-way ANOVA followed by Tukey–Kramer test).

**Extended Data Fig. 9.**
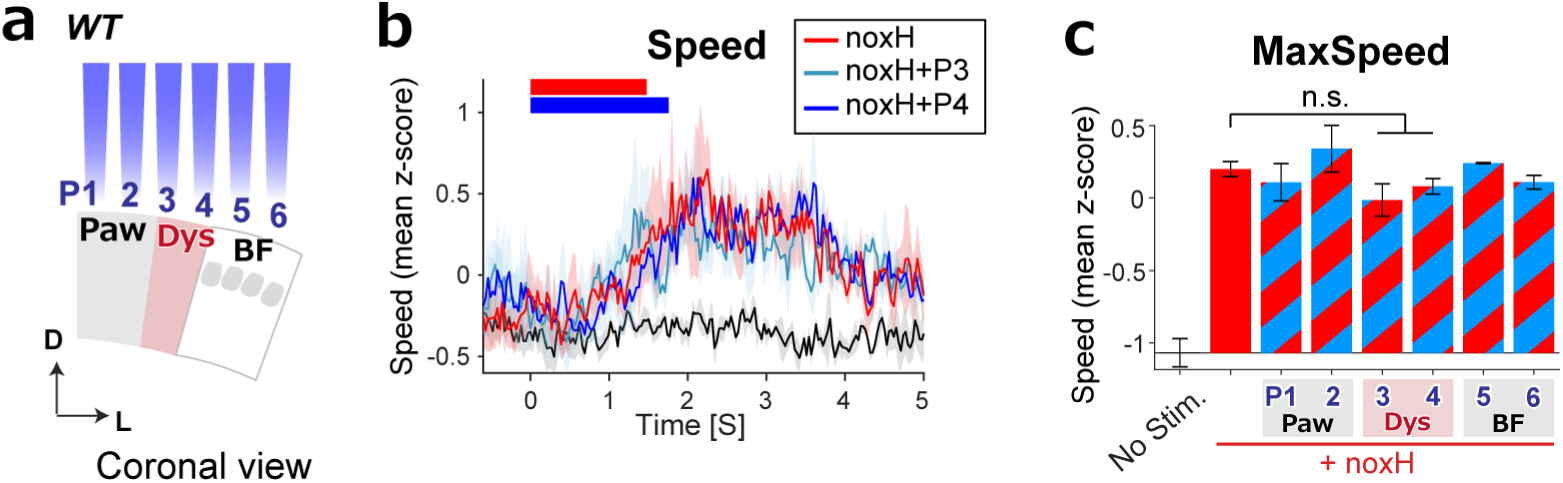
Blue light stimulation of Dys in control animals did not reduce the escape speed. **a**, Same setup as shown in Fig. 4e but in ChR2-negative wild-type (WT) animals. **b**, Blue light stimulations of P3 and 4 did not reduce the escape speed in WT mice (*n* = 3). **c**, Blue light stimulation at any position did not affect MaxSpeed (one-way ANOVA followed by Tukey– Kramer test).

**Extended Data Fig. 10.**
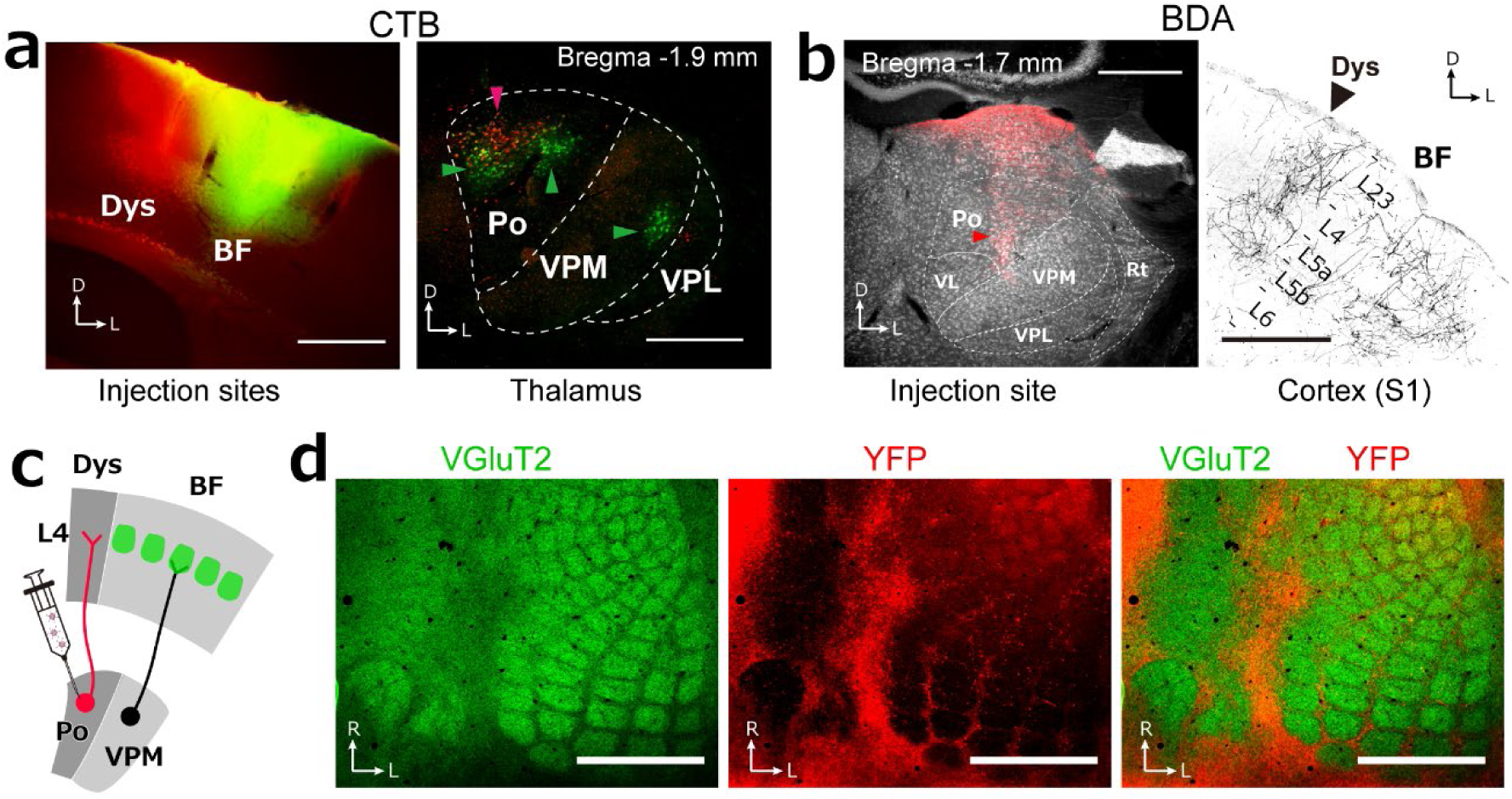
Dys receives input from the posterior medial thalamic nucleus. **a**, Retrograde labeling of posterior medial thalamic nucleus (Po) neurons after injection of Alexa Fluor 555- and 488-labelled cholera toxin subunit B into Dys (red) and BF (green), respectively. *Left*, Injection sites. *Right*, Somas of neurons projecting to Dys were observed in Po. The somas of neurons projecting to BF were observed in the ventral posterior medial nucleus (VPM) and Po. VPL, ventral posterior lateral nucleus. Scale bars, 500 μm. **b**, *Left*, Injection site of biotinylated dextran amine (BDA), an anterograde tracer. *Right*, Axon terminals from neurons in Po. VL, ventral lateral nucleus; Rt, reticular nucleus. Scale bars, 500 μm. **c**, Schema of AAV-DJ-YFP injection. **d**, Tangential section of S1 L4 co-labelled with anti-vGluT2 antibody (green) and axon terminals from Po (red). Scale bars, 1 mm. See also other tracer studies ^1,2^.

**Extended Data Fig. 11.**
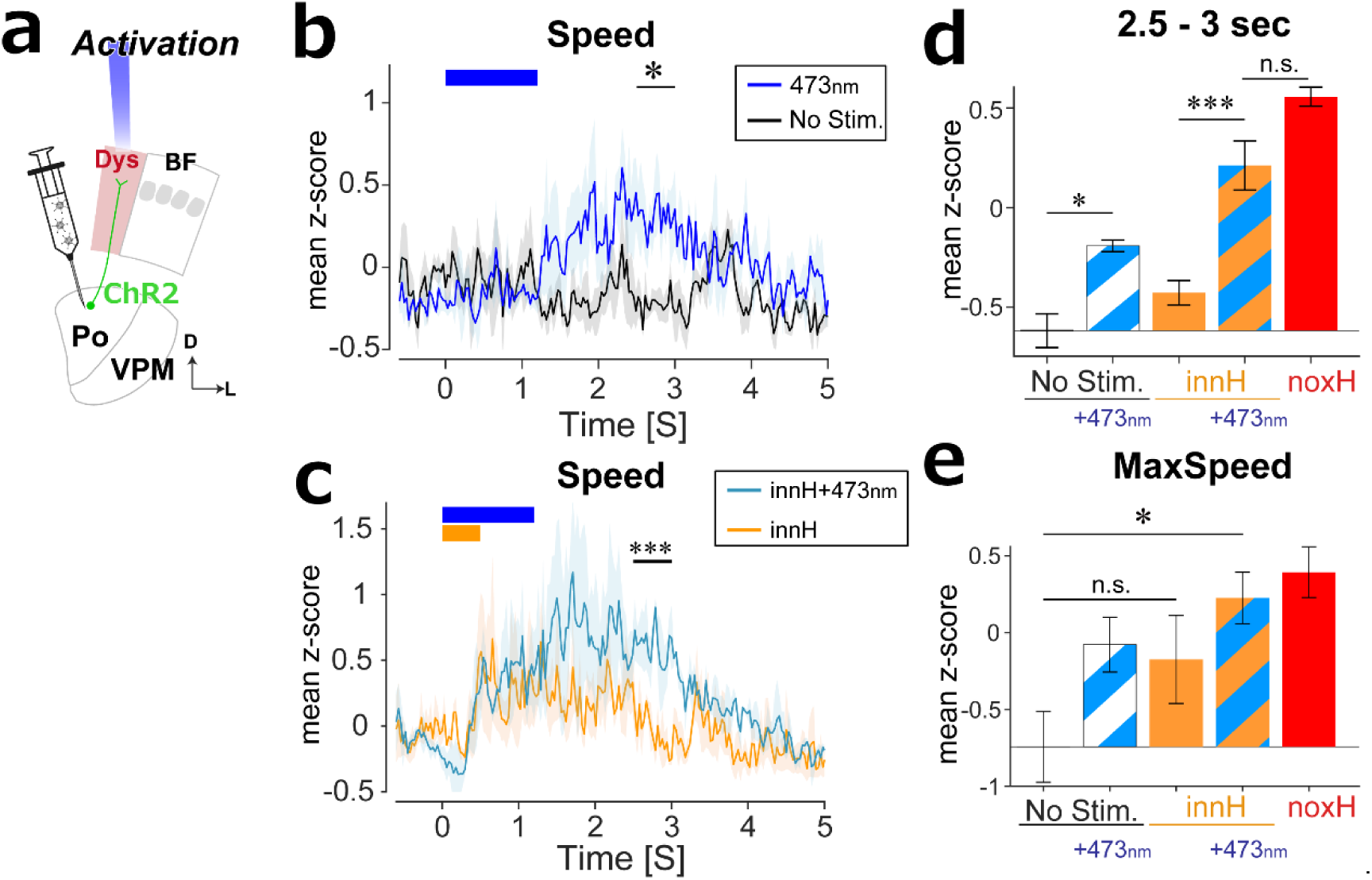
Optogenetical activation of thalamocortical fibers projecting to Dys induced escape responses. **a**, Same setup as shown in Fig. 4h. **b**, Average speed profiles with (473 nm, blue) and without (No Stim, black) optogenetic activation (*n* = 3 mice). **c**, Average speed profiles under innH condition with (innH + 473nm, cyan) and without (innH, orange) optogenetic activation **d**, Mean z-scores of the speed at 2.5–3 s. **e**, Means z-scores of MaxSpeed. n.s., not significant; \**P* < 0.05, \*\*\**P* < 0.001 by one-way ANOVA followed by Tukey–Kramer test.

**Extended Data Fig. 12.**
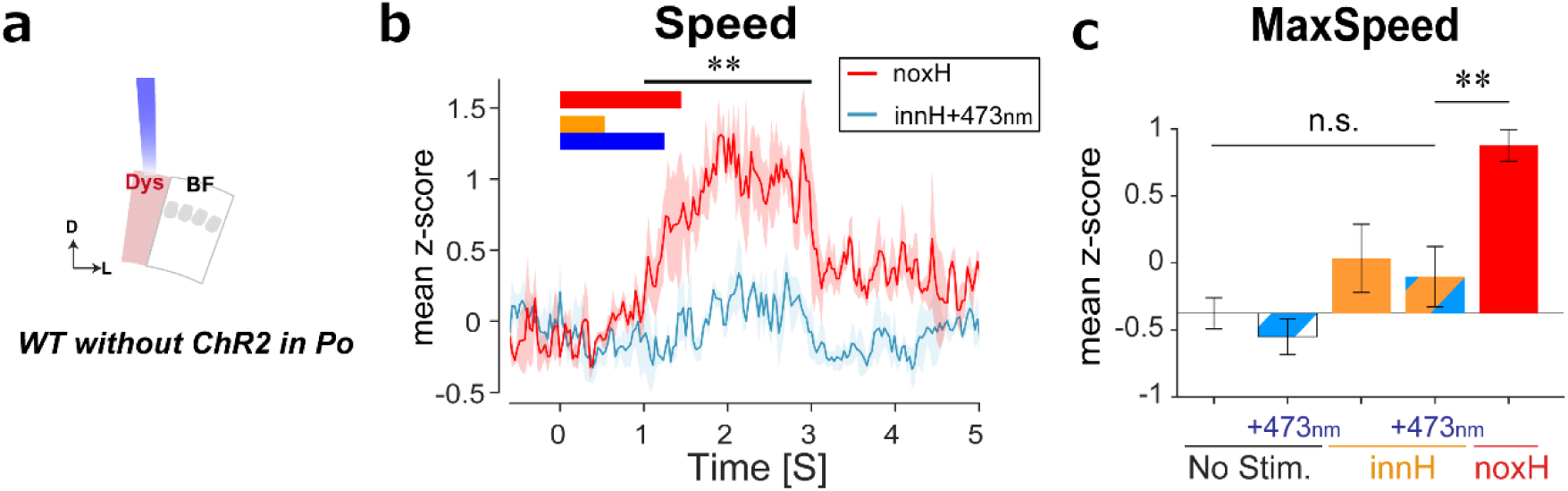
Blue light stimulation itself does not affect escape speed in animals not expressing ChR2. **a,** Same setup as the behavioral experiment in Fig. 4h in WT mice without AAV-ChR2 injection into Po. **b,** Mean z-scores of the speed profiles under the noxH condition and innH + 473 nm condition. Shading indicates SEM over three mice. **c,** Mean z-scores for maximum speed. The blue light (473 nm) combined with/without IR stimulation did not increase MaxSpeed. n.s., not significant; \*\**P* < 0.01 by one-way ANOVA followed by Tukey–Kramer test (*n* = 3).

